# A genome wide CRISPR screen reveals that HOXA9 promotes Enzalutamide resistance in prostate cancer

**DOI:** 10.1101/2023.12.15.571833

**Authors:** Michael V. Roes, Frederick A. Dick

## Abstract

Androgen receptor inhibitors are commonly used for prostate cancer treatment, but acquired resistance is a significant problem. Co-deletion of RB and p53 is common in castration resistant prostate cancers, however they are difficult to target pharmacologically. To comprehensively identify gene loss events that contribute to enzalutamide response, we performed a genome-wide CRISPR knockout screen in LNCaP prostate cancer cells. This revealed novel genes implicated in resistance that are largely unstudied. Gene loss events that confer enzalutamide sensitivity are enriched for GSEA categories related to stem cell and epigenetic regulation. We investigated the myeloid lineage stem cell factor HOXA9 as a candidate gene whose loss promotes sensitivity to enzalutamide. Cancer genomic data reveals that HOXA9 overexpression correlates with poor prognosis and characteristics of advanced prostate cancer. In cell culture, HOXA9 depletion sensitizes cells to enzalutamide, whereas overexpression drives enzalutamide resistance. Combination of the HOXA9 inhibitor DB818 with enzalutamide demonstrates synergy. This demonstrates the utility of our CRISPR screen data in discovering new approaches for treating enzalutamide resistant prostate cancer.

## Introduction

Prostate cancer is one of the most commonly diagnosed cancers among men and a leading cause of cancer related deaths (1). In the majority of cases, prostate cancer is an androgen dependent disease that is driven through the androgen receptor signaling pathway (2–4). First line therapy for recurrent prostate cancer patients acts to deplete hormones through androgen deprivation (3). However, some patients develop castration resistant prostate cancer, in which disease progresses in a low androgen environment (5). Second-generation AR inhibitors, such as enzalutamide, have led to improved patient outcomes for castration resistant prostate cancer (6, 7). However, these benefits are often short lived as patients develop resistance and enter a fatal stage (8).

An increasingly recognized mechanism of resistance to enzalutamide in castration resistant disease occurs through lineage plasticity or acquisition of a neuroendocrine fate (8–11). In this resistance scenario, an adenocarcinoma alters its lineage under the selective pressure of enzalutamide to a new fate that no longer expresses or needs the androgen receptor for proliferation (8). Treatment induced neuroendocrine prostate cancer is a clinically aggressive disease characterized by RB and p53 loss, low AR signaling and increased expression of neuroendocrine markers such as Chromogranin A (CHGA), Synaptophysin (SYP) and Neuron Specific Enolase (NSE)(8, 11). Quite commonly seen in clinical biopsies are mixed adenocarcinoma and neuroendocrine tumors that is indicative of lineage plasticity where cells are uncommitted a specific lineage (12). These patients are often treated with platinum-based chemotherapies but have dismal survival rates of about 24 months (13). This has led to an intense search for drivers of treatment induced neuroendocrine prostate cancer that has identified MYCN (14), SOX2 (15), SOX9 (16), FOXA1 (17), ONECUT2 (18), BRN2 (19), and PEG10 (20) among others. Unfortunately, only N-Myc regulation of EZH2 has thus far presented a therapeutic strategy to combat this advanced form of prostate cancer (10).

The HOX cluster transcription factor HOXA9 is known for its role in maintaining myeloid progenitor cells (21), and as an oncogene in acute myeloid leukemia where it is overexpressed in greater than 50% of cases (22). Known HOXA9 target genes such as ERG are known to contribute to prostate cancer as a target for recurrent chromosome translocations (23). In addition, HOX cluster deregulation and epigenetic alterations have been suggested to drive prostate cancer disease progression (24, 25). However, a role for HOXA9 in therapeutic resistance has yet to be reported.

In the current study we undertook a genome-wide CRISPR knockout screen in LNCaP prostate cancer cells to identify gene loss events that confer resistance or sensitivity to enzalutamide. This revealed a role for genes involved in epigenetic regulation and stem cell biology as key contributors to enzalutamide resistance. This screen identified HOXA9 whose loss promotes sensitivity to enzalutamide and whose overexpression drives resistance. Our analysis of prostate cancer patient genomic data reveals that HOXA9 copy number and expression are positively correlated with neuroendocrine features and a poor disease prognosis. Lastly, we find that experimental HOXA9 inhibitors can act in synergy with enzalutamide, and to restore responsiveness to enzalutamide in RB-p53 double mutant cells. This suggests that HOXA9 inhibition may represent an approach to treating enzalutamide-resistant prostate cancer patients.

## Results

### A Genome-wide CRISPR knock out screen implicates chromatin regulation and a stemness phenotype in enzalutamide (EZ) sensitivity

To investigate the landscape of genetic determinants to enzalutamide response and resistance in an unbiased manner, we performed a genome-wide CRISPR knockout (KO) screen in LNCaP prostate cancer cells (Fig. 1A). LNCaP cells were transduced with a lentiviral library encoding Cas9 and gRNAs targeting over 18,000 protein coding genes (26). The resulting pool of KO cells were split into 3 treatment and control replicates and subsequently treated with enzalutamide or DMSO. Cells were cultured continuously under treatment conditions for approximately two months. Next generation sequencing of guide RNA (gRNA) coding sequences from the initial and final timepoints for each replicate were used to identify genes whose presumptive mutations were enriched or de-enriched using MAGeCK (27).

**Figure 1:**
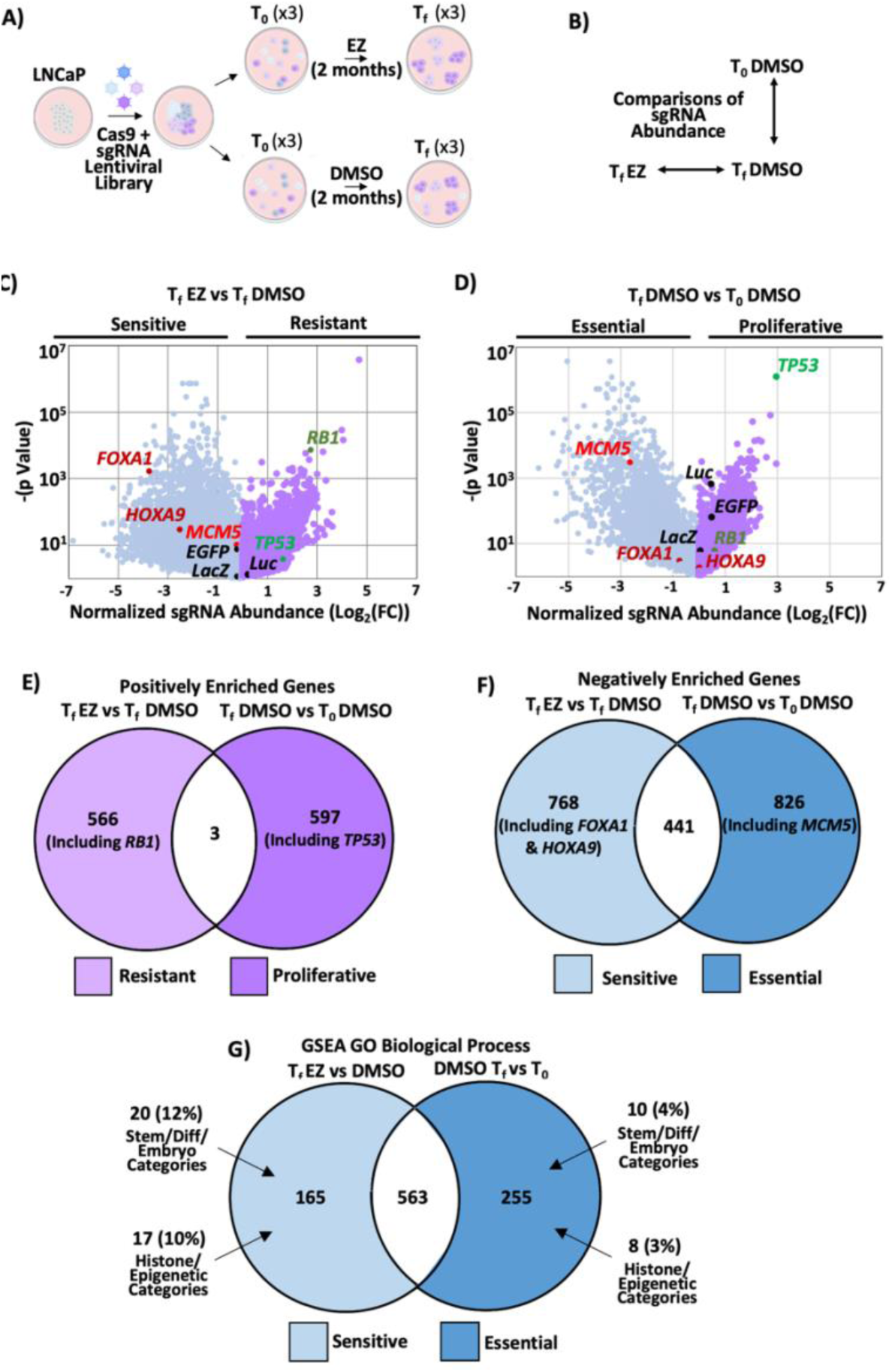
Genome-wide CRISPR knock out screen identifies stemness and epigenetic related genes as important for cell viability under enzalutamide treatment. **A**) Schematic of genome-wide CRISPR knockout screen workflow. LNCaP cells were transduced with a lentivirus pooled sgRNA library targeting over 18,000 protein coding genes. Infected cells were divided into 3 treatment and 3 control replicates and treated with enzalutamide or DMSO for approximately 2 months. **B**) Genomic DNA was extracted from cells from initial (T_0_) and final timepoints (T_f_), sgRNA sequences were PCR amplified and sequenced. MAGEK was used to compare sgRNA sequence abundance across treatments and timepoints. **C**) Volcano plot of sgRNA abundance scored for each gene in enzalutamide treated cells relative to DMSO at T_f_. Genes diminished in this analysis are labeled as ‘sensitive’ and illustrated with grey spots, while genes enriched in this comparison are labeled as ‘resistant’ in purple. Select genes or controls are annotated. **D**) Volcano plot of sgRNA abundance scored for each gene at the T_f_ and T_0_ timepoints in DMSO vehicle treated cells. **E**) Venn Diagram comparing overlap and number of genes positively enriched in the resistant and proliferative categories. All genes considered have a Log_2_ fold change > 1, *p* value < 0.05. Representative genes from each set are labeled. **F**) Venn Diagram of number of genes negatively enriched genes from the screen (Log_2_ fold change < -1, *p* value < 0.05). Representative genes from each set are shown. **G**) Venn diagram with number of GO processes found within each negatively enriched gene set annotated. GO terms related to stemness, differentiation or embryos (grey) and histones or epigenetics (green) are highlighted within the Enzalutamide Sensitive region.

We undertook a two-way analysis (Fig. 1B) in which gene enrichment or loss, both temporally and between treatment groups, allowed for inference of gene function in response to enzalutamide. This approach facilitated quality control of the screen workflow, as well as discovery of new genes that impacted the cellular response to enzalutamide. Genes enriched, or lost, at T_f_ in enzalutamide treatment compared with T_f_ DMSO control cells may indicate genes that confer enzalutamide sensitivity or resistance (Fig. 1C). The loss of *FOXA1* mutant cells in response to enzalutamide treatment in this experiment is consistent with its role as a driver of resistance in advanced prostate cancer (17, 28, 29). Similarly, the increased presence of *RB1* and *TP53* mutant cells is consistent with their loss of function in enzalutamide resistant disease (15, 28). However, gene loss events that score as sensitive, may reflect essential roles in cell physiology irrespective of drug treatment. In addition, loss of function mutations in negative growth regulators may simply increase proliferation in prolonged culture creating a false enzalutamide resistance phenotype. Comparison of T_f_ DMSO with T_0_ DMSO (Fig. 1D) was made to control for essential gene functions or roles in cell proliferation. Most non-targeting sgRNA controls directed against non-mammalian genes such a Luc, EGFP, and LacZ appear close to a Log_2_ fold change of zero and often lack statistical significance (Fig. 1D), consistent with a neutral effect on proliferation or survival over time. In the analysis of DMSO treated controls (Fig. 1D), loss of the MCM5 gene confirms its essential role in the proliferation of these cells and suggests its loss in enzalutamide treatment (Fig. 1C) isn’t related to drug action. As further support for our characterization of gene essentially, we identified all of the top 10 preferentially essential genes for LNCaP cells highlighted in the DepMap database, as de-enriched in our T_f_ DMSO vs T_0_ DMSO analysis (Supp. Table 1) (30). Analogously, *TP53* depletion is enriched over time in control culture (Fig. 1D), suggesting that cells deficient for p53 have a proliferative advantage that may contribute to its apparent resistance phenotype during enzalutamide selection (Fig. 1C). *RB1* deletion has little impact on overall proliferation over two months in control culture conditions (Fig. 1D), indicating its loss in response to drug treatment (Fig. 1C) represents a role in acquisition of enzalutamide resistance in this experiment. The accurate classification of these genes suggested our screen data is ideal for discovering new mechanisms of enzalutamide treatment resistance.

To search for novel genes implicated in enzalutamide response, we considered genes with Log_2_ Fold Change >1 or <-1, and with a p-value < 0.05. We compared gene lists created from the enzalutamide resistance analysis in Fig. 1C with proliferative genes in Fig. 1D. A Venn diagram illustrates the overlap of these gene sets in Fig. 1E and highlights the locations of genes used as controls in our screen. Gene sets in Fig. 1E show very little overlap between resistance genes and proliferative genes. This analysis identified 566 genes whose loss of function confers resistance to enzalutamide. Among negatively enriched genes with a Log_2_ Fold Change <-1, we compared gene lists from the enzalutamide sensitive category (Fig. 1C) with the essential gene category from DMSO cultured controls (Fig. 1D) as illustrated in Fig. 1F. This eliminated 441 genes that were shared between these lists and identified 768 genes in the sensitivity category for future study.

We performed gene set enrichment analysis (GSEA) on sensitivity and resistance gene lists to identify gene ontology (GO) biological processes or other relevant gene characteristics to better explain how LNCaP cells respond to enzalutamide (31). Analysis of the resistance gene set demonstrated an enrichment of a small assortment of functional categories, some of which are related to action potentials, T Helper 17 cell lineage, and defense against gram negative bacterium (Figure 1G). This seemingly random set of biological process terms is most likely because many genes in the resistance list are relatively unstudied and therefore lack categorization of a clear, well-defined function. Many possess gene names based on predicted primary amino acid structure and their homology to others. These include members of the ZNF, TMEM and RNF families of genes, among others. As such, their similarity to previously studied family members allow them to be categorized based on their DNA binding domains, transmembrane domains, ubiquitin ligase homology, respectively. In an effort to further elucidate putative drivers of enzalutamide resistance, we performed network and clustering analysis on enzalutamide resistance genes. As highlighted in Supplementary Figure 1A we noted one gene cluster with central nodes including ADRBK1, RHOA and PRCKA. In line with known functions of these genes and others within this cluster, we identified gene ontology terms enriched for this cluster related to G-protein coupled receptor, RAP1 and Hippo signaling pathways (Supp. Table 2). These results suggest that dis-regulation of these pathways through loss of ADRBK1, RHOA or PRCKA may mediate enzalutamide resistance. Overall, this analysis suggests acquired resistance to enzalutamide, through loss of function, represents a relatively unexplored area.

A GSEA analysis of sensitivity genes yielded 165 GO functional categories. Notably, 12% of these terms are related to stemness, differentiation or embryonic development and 10% related to histones or epigenetic regulation out of 165 processes significantly enriched in the enzalutamide sensitive category (Figure 1H, Supp. Table 3). This is striking, as significantly enriched GO terms amongst essential genes only yielded 3% and 4% of terms related to stemness and epigenetics, respectively. These results suggest that stemness and epigenetic pathways are important for LNCaP cell growth under enzalutamide treatment conditions, and that loss of these genes promotes sensitivity.

To identify new genes that contribute to enzalutamide resistance and drive prostate cancer progression we searched for sensitivity genes that are putative new drivers of resistance. HOXA9, a transcription factor known to function in embryonic development and differentiation of myeloid progenitor cells (22), was also identified in this category because its loss of function sensitizes cells to enzalutamide (Fig. 1C) and its loss of function in control culture conditions has no significant effect on viability (Fig. 1D). Furthermore, based on previously performed viability screens in DepMap, HOXA9 has a gene effect score of –0.15 and is not considered a dependent gene in LNCaP cells as highlighted in (Supp. Table 4) (30). Importantly, only 51 of 1805 screens uploaded to the database score HOXA9 as a dependent gene (Supp. Fig. 1B), further supporting that HOXA9 is not an essential gene and its effects in this screen are mainly related to enzalutamide response. HOXA9 was investigated further in this report based on these screen results and characteristics identified in subsequent experiments.

### RB and p53 deficient cells overexpress HOXA9 and are enzalutamide resistant

Co-deletion of RB and p53 is frequently seen in castration resistant and neuroendocrine prostate cancers (8). To model and characterize this advanced disease state *in vitro* we generated RB-p53 double knockout (DKO) LNCaP cells using CRISPR Cas9 (Fig. 2A). Validation of enzalutamide resistance was subsequently performed in short term (6 day) and long term (4 week) enzalutamide treatment settings on pools and isolated clones of DKO cells. DKO pools were treated with increasing concentrations of enzalutamide over an acute 6-day period and viability was determined with Alamar Blue staining. This revealed a moderately increased IC_50_ value in one of the two DKO pools compared with LNCaP parental cells (Fig. 2B). However, when cells were seeded at low density and continuously treated with enzalutamide over 4 weeks, only DKO pools formed visible colonies after staining with crystal violet (Fig. 2C,D). These results suggest that RB-p53 deficiency allows for the acquisition and selection of enzalutamide resistant clonal populations over prolonged treatment periods. Importantly, individual DKO clones also demonstrated colony forming ability in prolonged enzalutamide treatment (Supp. Fig. 2A and B) and were used for further study.

**Figure 2:**
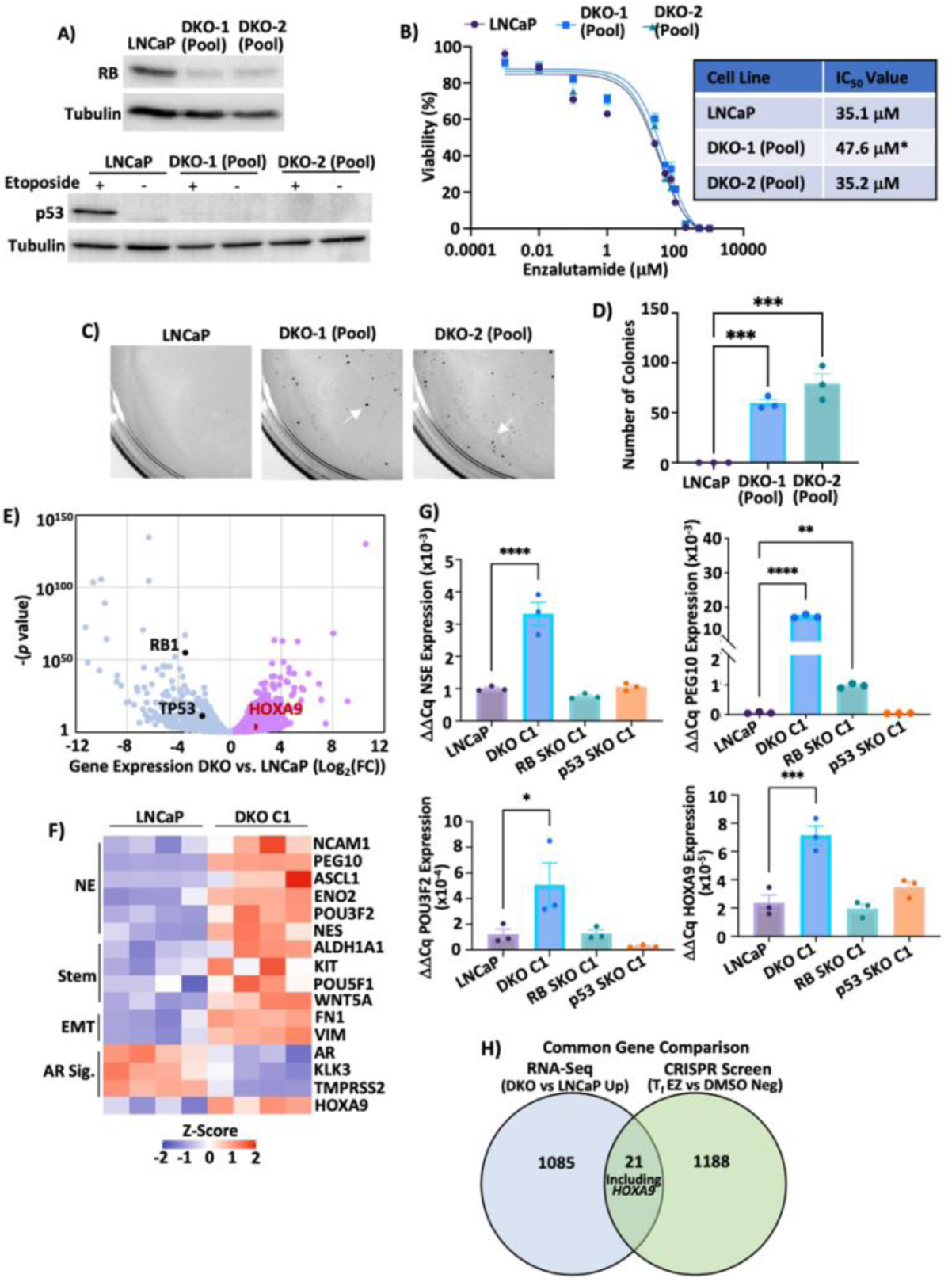
RB-p53 DKO cells are resistant to enzalutamide (EZ) and up-regulate stemness genes, such as HOXA9. **A)** Upper: Representative western blot highlighting CRISPR-Cas9 induced knockout of RB in RB-p53 DKO Pools. Cell lysates from LNCaP WT and RB-p53 DKO Pools probed for RB. Lower: Representative western blot highlighting CRISPR-Cas9 induced knockout of p53 in RB-p53 DKO Pools. Cell lysates from LNCaP WT and RB-p53 DKO Pools treated with etoposide or vehicle and probed for p53. **B)** Alamar Blue cell viability assay for LNCaP WT and DKO Pools treated with various concentrations of enzalutamide for 6 days. Representative non-linear regression line for each cell type is shown and generated by taking the mean viability values of each concentration from biological replicates. IC_50_ values were obtained by taking the mean best-fit IC_50_ value of biological replicates for each cell type and compared using one-way ANOVA. N = 5 or 6 biological replicates. **C)** Representative images from colony forming assay of LNCaP WT and DKO Pools. White arrows point to representative colonies stained with crystal violet. **D)** Analysis of colony forming assay from C. Number of colonies were counted using ImageJ software. **E)** Volcano plot of genes differentially expressed in RNA-seq analysis of DKO cells relative to LNCaP WT. Relative expression of negative controls RB1 and TP53 are labelled and identified in black. Relative expression of HOXA9 is labelled and identified in red. **F)** RNA-seq expression heat map of representative neuroendocrine (NE), stemness (Stem), epithelial-to-mesenchymal transition (EMT) and androgen receptor signature (AR Sig.) genes significantly and differentially expressed in DKO cells relative to LNCaP WT (n = 4 replicates). **G)** Comparative mRNA expression of labelled NE genes and HOXA9 in LNCaP WT, RB and p53 SKO and DKO clone C1 measured by RT-PCR. Bars show ΔΔCq values (mean ± SEM, n = 3 replicates). One-way ANOVA. **H)** Venn Diagram comparing overlap of genes significantly up-regulated in DKO cells (Log_2_ Fold Change > 1, p < 0.05) and negatively enriched in the CRISPR screen at the final timepoint (Log_2_ Fold Change < -1, p < 0.05). Where applicable: ****p < 0.0001, ***p < 0.001, **p < 0.01, *p < 0.05, ND – Not Detected, ns – Not Significant.

RNA sequencing was performed on DKO clone 1 and parental LNCaP cells to identify differentially expressed genes between these two cell lines (Fig. 2E). We identified up-regulation of neuroendocrine, stemness, and epithelial-to-mesenchymal transition genes, and down-regulation of androgen receptor target genes in DKO cells compared with LNCaP (Fig. 2F). These transcriptional changes were only seen in DKO cells, as single knockout clones of RB or p53, do not show up-regulation of neuroendocrine markers NSE, PEG10 and POU3F2 and fail to form colonies over a chronic enzalutamide treatment (Fig. 2G, Supp. Fig. 2C-D). Overall, these results suggest the DKO cells model the phenotypic and molecular features of enzalutamide resistant prostate cancer, and they demonstrate increased expression of HOXA9 (Fig. 2F,G).

We compared lists of significantly up-regulated genes in DKO cells and those de-enriched in the CRISPR screen (sensitivity genes). As highlighted in Fig. 2H and Supplementary Table 5 there were only 21 genes in common and this included HOXA9. Overall, these results suggest that the expression of the transcription factor HOXA9 may contribute to the acquisition of enzalutamide resistance.

### HOXA9 amplification is associated with poor overall survival, castration resistant disease, and neuroendocrine histology

Our CRISPR screen data suggests that HOXA9 is a potential mediator of enzalutamide resistance. We sought to extend these findings by analyzing publicly available genomic and transcriptomic datasets from prostate cancer patients using cBioPortal (32). We observed that increased HOXA9 copy number is correlated with shorter survival (Fig. 3A), where median survival was reduced from 132 months to 109. Since HOXA9 is up-regulated in enzalutamide resistant DKO cells, we investigated HOXA9 copy number in studies from hormone sensitive and castration resistant disease. HOXA9 was amplified, or exhibited single copy gain, in 39.3% of castration resistant cases compared with only 17.6% of patients in studies of hormone sensitive prostate cancer. Furthermore, in the MSKCC/DFCI 2018 dataset tumor samples are categorized as biopsies from primary or metastatic sites (33). Genomic gain of HOXA9 was found in 40% of metastatic tumors compared with 17% in primary prostate tumors (Figure 3D), showing that increased HOXA9 copy number is associated with advanced prostate cancer. Lastly, we analyzed the SU2C/PCF Dream Team 2019 study as it relates genomic, transcriptomic, and histologic data from castration resistant patients (28). Fig. 3D shows that HOXA9 is genetically altered or mis-expressed in 10% of samples. Strikingly, HOXA9 amplifications and over-expression accounts for 95% of these alterations, further suggesting that HOXA9 has an oncogenic role in advanced prostate cancer. We next evaluated HOXA9 mRNA expression in samples with histologic neuroendocrine features and they are correlated with higher HOXA9 expression (Fig. 3E). Furthermore, when categorized based on histopathologic classification, HOXA9 expression is highest in tumors with a mixed adenocarcinoma and neuroendocrine histology, compared with pure adenocarcinoma or small-cell features (Fig. 3F). Lastly, since RB loss was key to discovering HOXA9 upregulation, we compared RB1 and HOXA9 expression in samples from this study. We found an inverse correlation between RB1 and HOXA9 expression (Fig. 3G). Overall, these clinical genomic observations indicate that increased HOXA9 expression is associated with advanced prostate cancer and may be an important mediator in the cellular response to enzalutamide.

**Figure 3:**
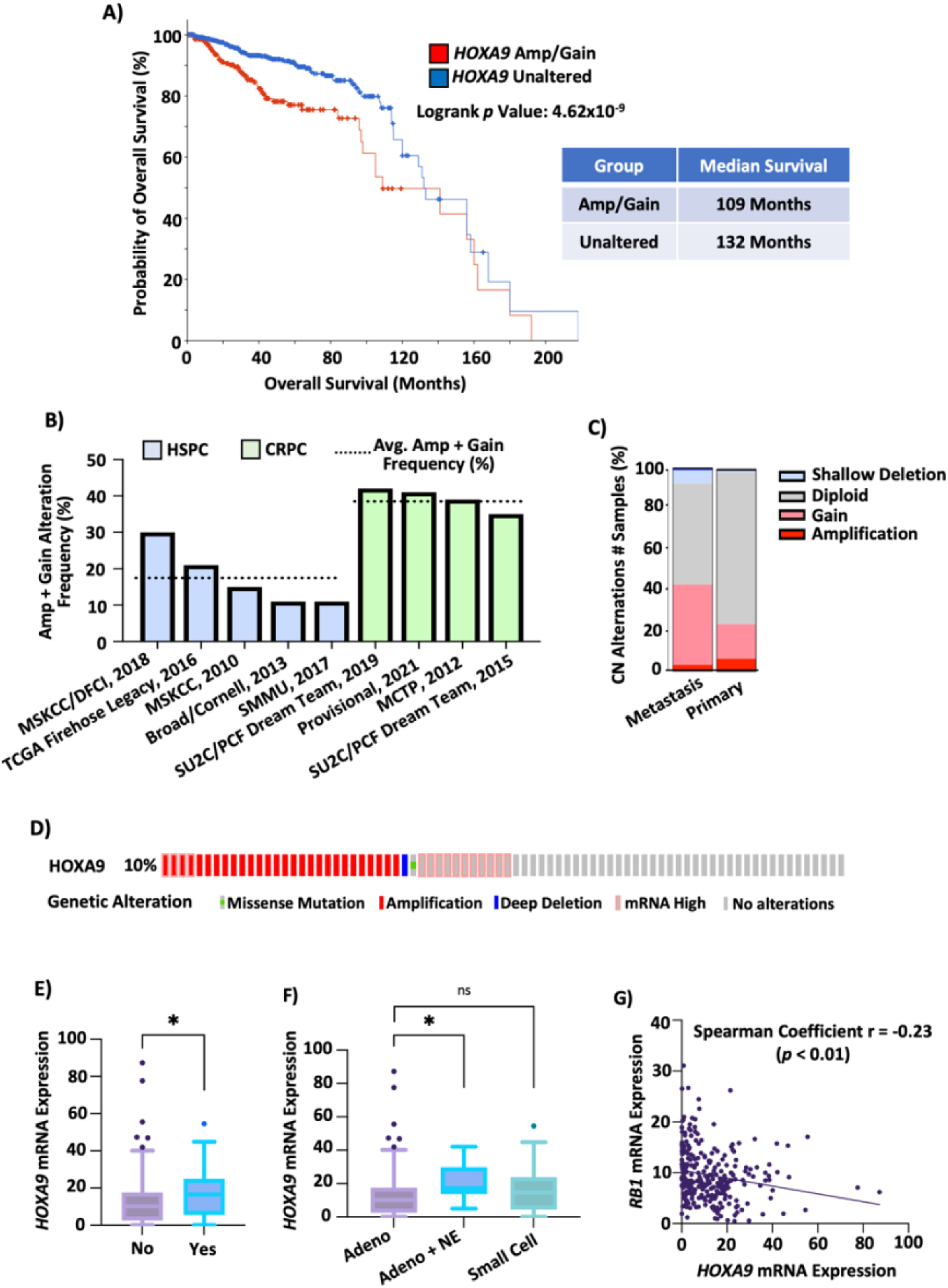
HOXA9 amplification and gain in prostate cancer clinical samples is associated poorer overall survival, an advanced diseased state and neuroendocrine features. **A)** Kaplan Meir curve comparing overall survival of prostate cancer patients with or without HOXA9 genomic amplification or gains across all prostate cancer studies in cBioPortal. Median overall survival for each cohort is shown. Log Rank Test performed. **B)** Frequency of HOXA9 amplification + gains across hormone sensitive prostate cancer (HSPC) and castration resistant prostate cancer (CRPC) datasets from cBioPortal. Average frequency for cohorts is shown. Welch’s unpaired t-test performed. **C)** Genomic alterations within primary and metastatic prostate tumour samples from the MSK/DFCI, Nature Genetics 2018 HSPC dataset. **D)** Oncoprint for HOXA9 displaying genomic and transcriptomic alterations in CRPC tumour samples from the SU2C/PCF Dream Team, PNAS 2019 dataset. **E)** Tukey plots of HOXA9 mRNA expression (FPKM Poly A) in tumour samples with or without NE features from CRPC dataset in D). Welch’s one-tailed t test. **F)** Tukey plots of HOXA9 mRNA expression across pathology classifications in CRPC dataset from D). Welch’s one-tailed t tests. **G)** Correlation between RB1 and HOXA9 mRNA expression in CRPC dataset from D). Simple linear regression line and Spearman Coefficient are shown. 2-sided t-test. Where applicable: ****p < 0.0001, ***p < 0.001, **p < 0.01, *p < 0.05, ns – Not Significant.

### HOXA9 overexpression drives enzalutamide resistance

Our genome-wide CRISPR knockout screen suggests that loss of HOXA9 sensitizes cells to enzalutamide. To validate these findings and further interrogate the role of HOXA9 in mediating enzalutamide response, we transduced cells with short-hairpin RNA (shRNA) to knock-down HOXA9 in LNCaP WT (LNCaP shHOXA9) and RB-p53 DKO cells DKO shHOXA9). Since HOXA9 protein levels are low in LNCaP and DKO cells, validation of our knockdown system was performed in HEK-293T cells that have higher HOXA9 expression levels (Supp. Fig 3A). We subsequently performed the same short term and long term enzalutamide resistance assays in LNCaP and DKO shHOXA9 cells as described above. Two distinct shRNAs against HOXA9 increased sensitivity of LNCaP cells to enzalutamide as determined by lower IC_50_ values compared with LNCaP expressing control shRNA (Fig. 4A). In contrast, the same HOXA9 directed shRNAs in DKO cells failed to sensitize cells to enzalutamide in a six day treatment period (Fig. 4B). However, when seeded at low density and treated with enzalutamide over four weeks, DKO shHOXA9 clones formed fewer colonies and smaller colonies compared with DKO control shRNA expressing cells (Fig. 4C,D, and Supp. Fig 3B). Collectively, these results further support our screen findings that HOXA9 loss of function sensitizes LNCaP cells to enzalutamide and this is seen in both LNCaP and enzalutamide resistant DKO cells.

**Figure 4:**
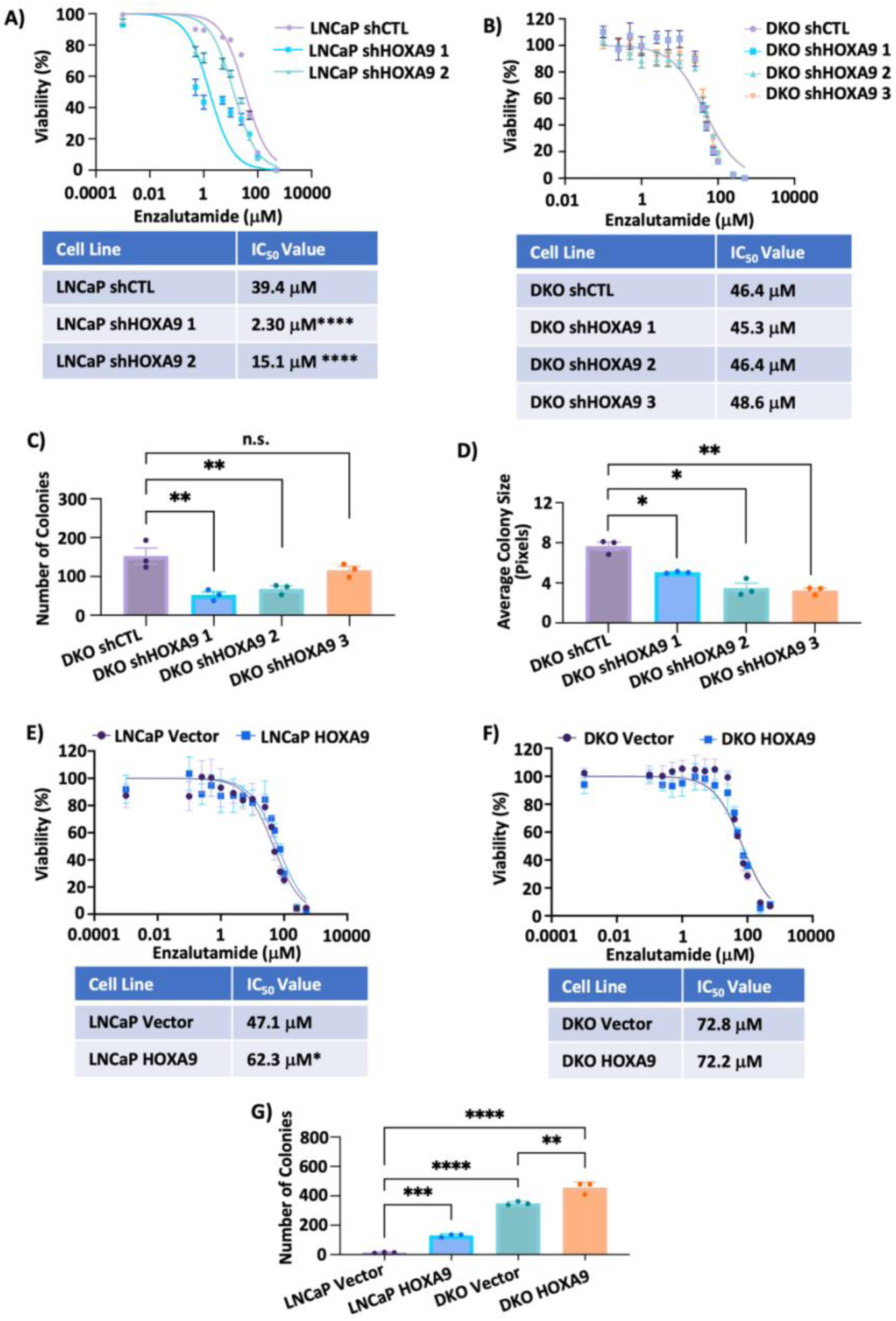
HOXA9 knockdown sensitizes cells to enzalutamide, whereas HOXA9 overexpression drives enzalutamide resistance. **A and B)** Alamar Blue cell viability assay for LNCaP (A) and DKO (B) shCTL and shHOXA9 clones treated with various concentrations of EZ for 6 days. Representative non-linear regression line for each cell type is shown and generated by taking the mean viability values of each concentration from biological replicates. IC_50_ values were obtained by taking the mean best-fit IC_50_ value of biological replicates for each cell type and compared using one-way ANOVA. N = 6 biological replicates. **C and D)** Number of colonies (C) and average colony size (D) in DKO shCTL and shHOXA9 cells following EZ colony forming assay. Colony number and size quantified using ImageJ software. One-way ANOVA. **E and F)** Alamar Blue cell viability assay for LNCaP (A) and DKO (B) HOXA9 overexpressing (HOXA9) or empty vector (Vector) control cells treated with various concentrations of EZ for 6 days. Representative non-linear regression lines and IC_50_ values obtained using method described above. Unpaired Welch’s t-test. N = 6 biological replicates. **G)** Number of colonies in LNCaP and DKO HOXA9 OE or EV cells following EZ colony forming assay. Colony number quantified using ImageJ software. One-way ANOVA. Where Applicable: ****p < 0.0001, ***p < 0.001, **p < 0.01, *p < 0.05, ns – Not Significant.

Since elevated HOXA9 copy number and expression levels are associated with castration resistance and poor overall survival in prostate cancer patients we investigated its overexpression. We transduced LNCaP and DKO cells with HOXA9 cDNA expression or empty vectors to generate overexpression lines (Supp. Fig 3C). In contrast to HOXA9 depletion, HOXA9 overexpression in LNCaP cells drove enzalutamide resistance, identified by an increased IC_50_ in short term treatment (Fig. 4E), and the formation of significantly more colonies over a four week treatment period (Fig. 4G, Supp. Fig. 3D). We also tested whether HOXA9 overexpression in DKO cells, that already up-regulate HOXA9, can further increase enzalutamide resistance. Although no difference was observed in IC_50_ value between DKO and HOXA9 overexpressing derivative cells (Fig. 4F), DKO HOXA9 over expressing cells formed significantly more colonies than either DKO or LNCaP HOXA9 overexpressing cells (Fig. 4G). These experiments demonstrated that HOXA9 overexpression leads to enzalutamide resistance, consistent with its elevated expression in castration resistant prostate cancer.

### HOXA9 inhibitors block expression of downstream targets and are synergistic with enzalutamide

Our findings that HOXA9 depletion sensitizes prostate cancer cells to enzalutamide suggests that therapeutic inhibition of HOXA9 in combination with enzalutamide could be a strategy for treating enzalutamide-resistant prostate cancer patients. We used an experimental HOXA9 inhibitor called DB818, that is known to inhibit binding of HOXA9 to DNA (34). FLT3 is a receptor tyrosine kinase that is known to be expressed in acute myelogenous leukemia in a HOXA9 dependent manner. We confirmed that FLT3 is also overexpressed in DKO cells compared to LNCaP by qRT-PCR (Fig. 5A). We treated LNCaP and DKO cells with increasing concentrations of DB818 and observed a dose dependent reduction in FLT3 expression in both LNCaP and DKO cells, at both the mRNA and protein levels (Figures 5B,C). Western blot analysis of LNCaP and DKO lysates similarly confirms that FLT3 expression is reduced by DB818 in a dose dependent manner.

**Figure 5:**
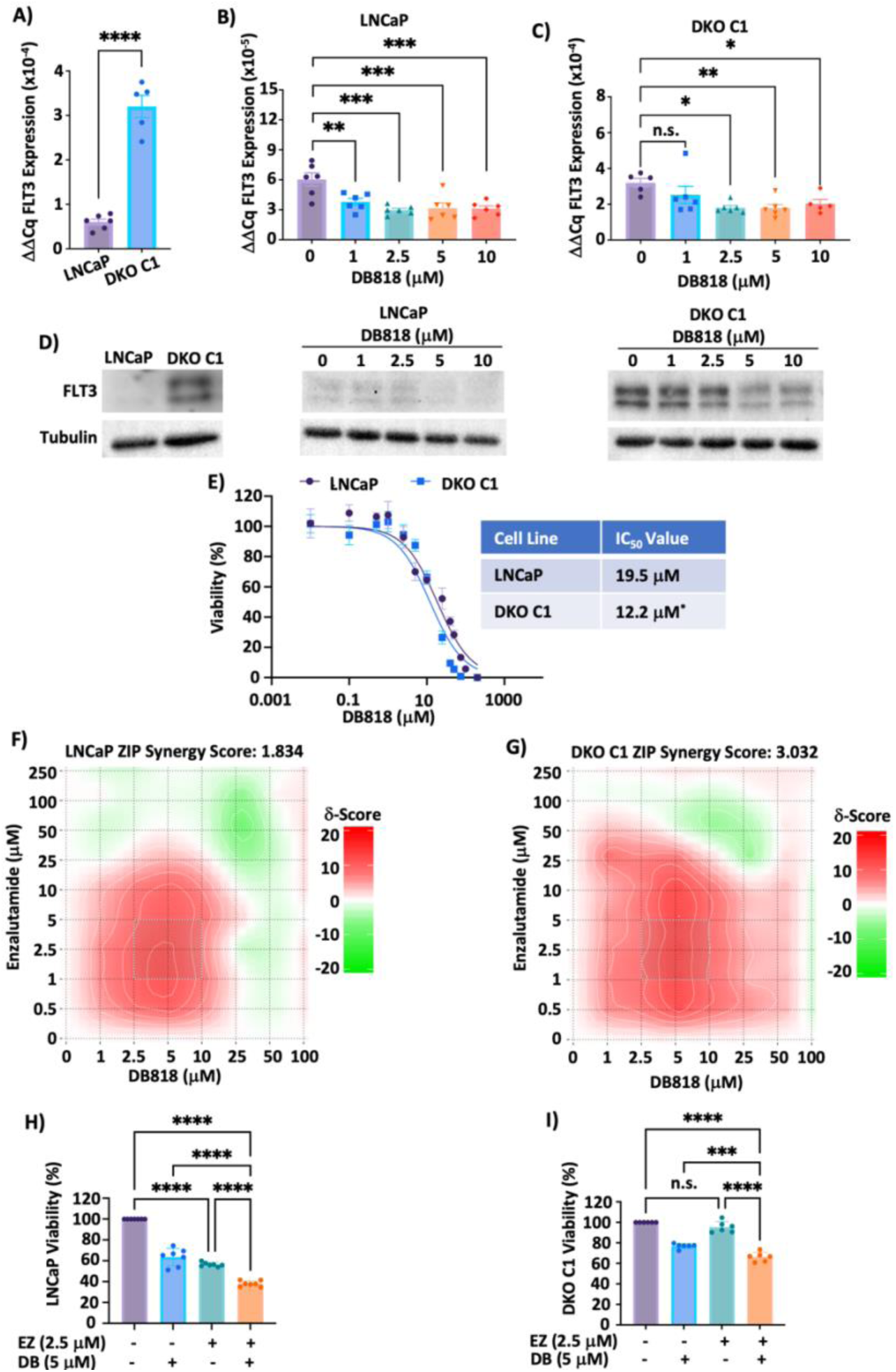
RB-p53 DKO cells are more sensitive to the HOXA9 inhibitor DB818 and have greater synergistic effects when treated in combination with enzalutamide. **A)** mRNA expression of HOXA9 target gene FLT3 in LNCaP WT and RB-p53 DKO-C1 cells measured by RT-PCR. Bars show ΔΔCq values (mean ± SEM, n = 3 replicates). Unpaired Welch’s t test. **B and C)** mRNA expression of FLT3 in LNCaP (B) and DKO-C1 (C) cells treated with various concentrations of the HOXA9 inhibitor DB818. **D)** Western blot of cell lysates from LNCaP WT and DKO-C1 treated as in A-C and probed for FLT3. **E)** Alamar Blue cell viability assay for LNCaP and DKO-C1 cells treated with various concentrations of DB818 for 6 days. Representative non-linear regression lines and IC_50_ values obtained using method described above. Unpaired Welch’s t-test. N = 6 biological replicates. **F and G)** Synergy maps for LNCaP (F) and DKO-C1 (G) cells treated with various combinations of DB818 and EZ. Synergistic (red) and antagonistic (green) dose regions are highlighted. The Most Synergistic Areas (MSA) for each cell type is shown as a dashed box on map. Calculated ZIP synergy scores for each cell type are also shown. **H and I)** Quantification of viability values from F and G when treated with single drug (EZ - enzalutamide or DB - DB818) or combination of both drugs at indicated doses in LNCaP (H) or DKO C1 (I) cells. N = 6 biological replicates. One-way ANOVA. Where applicable: ****p < 0.0001, ***p < 0.001, **p < 0.01, *p < 0.05, ns – Not Significant

LNCaP and DKO cells were treated with increasing concentrations of DB818 over a 6-day treatment period (Fig. 5E). This revealed that DKO cells are more sensitive to the HOXA9 inhibitor. Since DKO cells express higher levels of HOXA9 than LNCaP, this may explain their sensitivity to this agent and this motivated us to investigate its effects in combination with enzalutamide. We treated LNCaP and DKO cells with 80 different drug combinations for 6-days and determined viability using Alamar Blue staining. This data was used to generate the synergy plots in Fig. 5F,G. Red highlighted areas within the plots denote synergy between the 2 drug concentrations, whereas green highlighted areas denote antagonism. The greatest synergist area, marked by the dashed box in each plot, is found at low dose combinations for each cell type, ranging from 2.5-10 μM DB818 and 1-5 μM enzalutamide. This high level of synergy in LNCaP and DKO cells is further highlighted when quantifying cell viability following combination treatment with enzalutamide and DB818 compared with either drug alone, as in Fig. 5H,I. For example, LNCaP cells treated with 2.5 μM enzalutamide causes a significant reduction in cell viability, which is further reduced with the addition of 5 μM DB818 (Figure 5H). Importantly, in enzalutamide resistant DKO cells, cell viability is unaffected with 2.5 μM enzalutamide treatment, but is significantly reduced with the addition of 5 μM DB818 (Figure 5I). Furthermore, a greater overall ZIP synergy score in DKO cells of 3.002 compared with 1.834 in LNCaP cells further suggests that elevated HOXA9 expression may enhance DB818 effects to counteract enzalutamide resistance (Figure 5 F,G). Overall, these results suggest that the HOXA9 inhibition can sensitize cells to enzalutamide, particularly at low concentrations of both drugs, and may be of therapeutic benefit for treating enzalutamide-resistant prostate cancer patients.

## Discussion

This study generated a comprehensive map of gene roles in the cellular response to enzalutamide. We leveraged this data to identify previously unappreciated pathways in therapeutic response and validate HOXA9 as a driver of EZ resistance. We have shown that HOXA9 is up-regulated in the context of targeted RB and p53 deletion in culture experiments and it contributes to enzalutamide resistance. Analysis of prostate cancer genomic data indicates its expression correlates with poor outcomes, and its elevated expression occurs preferentially in advanced disease. Importantly, the HOXA9 inhibitor DB818 can act synergistically with enzalutamide, suggesting HOXA9 inhibition or it downstream targets may offer a new therapeutic approach to prostate cancer.

Genome-wide CRISPR knock out screens in which resistance and sensitivity to a targeted therapeutic agent such as enzalutamide are determined, provide unbiased information about gene dependencies in a cancer therapeutic context (35). In this study we categorized gene loss events into categories related to proliferation or gene essentiality, and sensitivity or resistance. Our analyses identified genes that have previously been shown to have roles in mediating enzalutamide resistance, such as RB1 and FOXA1 (15, 17, 29, 36), as well as known essential genes for proliferation in culture that have been identified from DepMap CRISPR screens in LNCaP cells (30). Our network and clustering analysis of resistance genes identified functions related to signal transduction, potentially through RAP1 and Hippo pathways. However, it is intriguing to note that many of the putative resistance genes we identified are relatively unexplored or have unknown functions. We considered if the two-month period in which we carried out our screen was sufficient to capture resistance genes. We observed that RB-p53 DKO cells form colonies after one month in enzalutamide containing media, suggesting they readily acquire resistance. In addition, our CRISPR screen readily identified RB in a two month period suggesting our screen conditions were robust for detecting resistance genes. We expect our screen will serve as a valuable resource for others studying the genetics of enzalutamide resistance particularly as this appears to be a largely unexplored area.

The physiological role of HOXA9 in the maintenance of hematopoietic stem cells has been well studied (37–39). Furthermore, HOXA9 knockout mice display decreased myeloid progenitor cells and elevated levels of erythroid cells. HOXA9 is overexpressed in 70% of acute myeloid leukemias (AML) and is considered the most important prognostic marker for AML patients (40, 41). We think there are a number of unappreciated similarities with castration resistant prostate cancer. Transdifferentiation or lineage plasticity suggests a role for a stem cell like state in the adenocarcinoma, neuroendocrine, and mixed lineage that is observed in advanced prostate cancer (12, 16). In this study we demonstrate that HOXA9 mRNA expression in castration resistant prostate cancer is highest in tumours with mixed histology compared with pure prostate adenocarcinoma or small cell carcinoma. Furthermore, in addition to upregulation of HOXA9, RB and p53 DKO cells also upregulate other stemness and EMT genes such as *SOX9*, *POU5F1*, *WNT5A* and *VIM* suggesting HOXA9 and its signaling pathway may represent a pharmacologic target within this enzalutamide resistant category. Our data supports the conclusion that HOXA9 upregulation is a previously unappreciated driver of enzalutamide resistance in advanced prostate cancer.

Currently, there are no HOXA9 inhibitors in clinical use for cancer treatment. However, the oncogenic role for HOXA9 in a number of cancer types, including AML, malignant glioblastoma and colorectal cancer suggests it and its downstream targets are an appealing potential therapeutic target (42–44). DB818 has been shown to suppress growth in 3 AML cell lines (44). The HOXA9 peptide inhibitor HXR9 and a derivative HTL-001 are effective in treating a variety of cancer cell and mouse models (45, 46). In this study we have shown that CRISPR-Cas9 KO, shRNA knockdown and small molecule inhibition of HOXA9 can all sensitize prostate cancer cells to enzalutamide. The synergy observed when co-treating LNCaP WT and RB-p53 DKO cells with enzalutamide and DB818 further suggests that inhibition of HOXA9 and its pathway may be a rational therapeutic approach for re-sensitizing enzalutamide-resistant prostate cancers. Overall, our systematic analysis of gene roles in enzalutamide response and resistance has revealed a new category of therapeutic targets to improve outcomes for prostate cancer patients.

## Methods

### Cell culture

LNCaP cells were obtained from Dr. Alison Allan at Western University and maintained in RPMI-1640 supplemented with 10% fetal bovine serum and 1% L-Glutamine-Penicillin-Streptomycin. HEK-293T cells were maintained in DMEM supplemented with 10% fetal bovine serum and 1% L-Glutamine-Penicillin-Streptomycin. LNCaP cells, CRISPR derived knock outs, and shRNA or HOXA9 overexpressing derivatives were verified by STR analysis using services from TCAG (Toronto).

### CRISPR screening

Toronto Knockout Version 3 library was purchased from Addgene (#90294) (26), and amplified and packaged as lentivirus as follows. HEK-293T cells were transfected at ∼50% confluency with the plasmid library, psPAX2 (Addgene # 12260) and pMD2.G (Addgene # 12259) in a 4:3:1 mass ratio with Lipofectamine 2000 (Invitrogen # 11668019) in a 2:1 ratio of Lipofectamine 2000: Plasmid DNA. Virus was collected 2 days after transfection, spun down at 1500 RPM for 10 minutes and the resulting supernatant was filtered through 0.45-micron filter. LNCaP cells were infected at an MOI of ∼0.3 with 8 μg/mL polybrene. Infected cells were selected with 2 μg/mL puromycin for approximately 2 weeks and subsequently divided into 6 pools (3 control and 3 treatment replicates). Pools were then passaged and treated with 75 μM enzalutamide (EZ) or equal volume DMSO for approximately 2 months. Cells were collected from each pool at the initial (passage immediately before EZ/DMSO treatment) and final timepoints (day 69) for genomic DNA extraction and PCR amplification of sgRNA sequences. The resulting sgRNA sequence amplicons from all 12 samples (3 control and 3 treatment groups at initial and final timepoints) were sequenced on an Illumina Next Seq instrument. Processed read data is available at GEO226977. Analysis for sgRNA abundance across treatments and timepoints was performed using MAGECK (27).

### CRISPR knockout, shRNA knockdown and cDNA over-expression cell line generation

To generate RB and p53 CRISPR-Cas9 single knockout cells (SKO), RB1 and TP53 sgRNA sequences highlighted in Supplementary Table 6 were cloned in a LentiCRISPRv2 (Addgene # 52961) backbone (47). Lentiviral production and infection were performed as described above. RB-p53 double knockout cells (DKO) were generated by subsequent infection of RB SKO cells. TP53 sgRNA sequences (TP53_1 and TP53_2 sgRNA sequences used to make RB-p53 DKO Pool 1 and Pool 2, respectively) were cloned into a pLX-sgRNA (Addgene # 50662) backbone (48). Lentiviral production and infection were performed as described above. Cells were selected using 3 ug/mL Blasticidin. All clones were generated by limiting dilutions. For generation of HOXA9 knockdown (KD) cells, shCTL and shHOXA9 constructs were purchased from Horizon Discovery (RHS4430-200210793, RHS4430-200285534, RHS4430-200285691). Short-hairpin sequences were subsequently excised using XhoI and MluI, and cloned into a pINDUCER11 miR RUG (Addgene # 44363) backbone (49). LNCaP WT, RB-p53 DKO Clone C1 cells and HEK 293T cells were infected with lentivirus containing these constructs as described above and subsequently treated with 500 ug/mL Zeocin for 2 weeks. For generation of HOXA9 overexpressing cells, pINDUCER21 (ORF-EG) and pINDUCER21-HOXA9 were purchased from Addgene (#46948 and #51302, respectively) (49, 50). LNCaP WT and RB-p53 DKO Clone C1 cells were infected with lentivirus containing these constructs (pINDUCER21 (ORF-EG) used for vector control and pINDUCER21-HOXA9 used for HOXA9 overexpression) as described above and subsequently FACS sorted using GFP fluorescence.

### Enzalutamide and HOXA9 inhibitor resistance assays

For acute resistance assays, cells were seeded into 96-well plates. After 3 days, cells were treated with various concentrations of DB818 and/or enzalutamide. Following a 6-day treatment period, media was replaced and Alamar blue cell viability dye (Invitrogen # DAL1100) was added at a concentration of 10%. After 3 hours of incubation cell viability was measured using a Synergy H4 hybrid reader (BioTek, USA) using excitation/emission wavelengths of 560nm/590nm. Fluorescence values were corrected using a blank of medium and Alamar blue, and then normalized to untreated cells. Normalized values for each biological replicate were fit to a curve using nonlinear regression ([Inhibitor] vs. Response – 3 parameters) or ([Inhibitor] vs. Normalized Response). IC_50_ values were obtained from the best-fit values and averaged across biological replicates for each cell type. Synergy plots and ZIP synergy scores were evaluated with Synergy Finder v3.0 using calculated percent viability values for each cell type as discussed above (51). For chronic enzalutamide, colony forming assays, cells were seeded at low density (10,000 cells/ 10 cm plate) and treated every 6 days with 50 μM EZ for approximately 1 month. Cells were subsequently fixed with methanol, stained with crystal violet, and imaged. Images were analyzed using ImageJ software to quantify colony number and size (52).

### Immuno-precipitation (IP) and Western blotting

Cells were grown to 60-80% confluency prior harvesting. For preparation of nuclear extracts and subsequent IP, cells were initially lysed with hypotonic lysis buffer (10 mM Tris pH 7.5, 10 mM KCl, 3 mM MgCl_2_, 1 mM EDTA) containing the following protease inhibitors: 1mM DTT, 1mM PMSF, 0.5 mM NaF, 0.1 mM Na_3_VO_4_, 5 μM Aprotinin and 5 μM Leupeptin. Nuclei were pelleted by and subsequently lysed using GSE buffer (20 mM Tris pH 7.5, 420 mM NaCl, 1.5 mM MgCl_2_, 0.2 mM EDTA, 25% Glycerol) containing 0.1% NP-40 and the following protease inhibitors: 2.5 mM DTT, 0.5 mM NaF, 0.2 mM Na_3_VO_4,_ 5 μM Aprotinin and 5 μM Leupeptin. Samples were standardized for protein concentration using the Qubit protein assay (Invitrogen #Q33211). 3 mg of protein from each sample were incubated with 3 ug of HOXA9 (Millipore # 07-178) or Rabbit IgG Isotype control (Invitrogen # 02-6102) antibodies overnight. Protein-antibody complexes were precipitated using 1.2 mg of magnetic Dynabeads Protein G (Invitrogen #10009) and subsequently denatured by diluting in SDS-PAGE sample buffer and boiling at 95°C for 5 minutes. For whole cell extracts, lysates were harvested with RIPA lysis buffer containing the following protease inhibitors: 1 mM DTT, 5 mM NaF, 0.25 mM Na_3_VO_4,_ 5 μM Aprotinin and 5 μM Leupeptin. For p53 immunoblots, cells were treated with 100 μM etoposide (Sigma # E1383) or equal volume DMSO (Fisher # BP231) for 8 hours prior to harvesting. Samples were standardized for protein concentration using the Qubit and denatured in Laemmli buffer to 1X and boiled at 95°C for 5 minutes. All protein samples were separated by SDS-PAGE and transferred on to PVDF membranes (Bio-Rad #1704274). Membranes were blocked with 5% skim-milk diluted in 1X TBST and incubated with primary antibodies overnight [RB (BD Pharmingen #554136, 1:500), p53 (Pierce #MA5-12453, 1:500), HOXA9 (Abcam # 140631, 1:1000), α-Tubulin (Cell Signaling Technology #2125, 1:1000). Blots were incubated with secondary antibodies [Rabbit (Cytiva #NA934V, 1:5000) or Mouse (Millipore #401215, 1:5000) and visualized using chemiluminescent substrate (SuperSignal West Dura Extended Duration Substrate #34075).

### Quantitative PCR and RNA Sequencing

Cells were grown to 60-80% confluency prior to harvesting. RNA was extracted from cells using Monarch Total RNA Miniprep Kit (NEB # T2010S). Any remaining DNA was degraded using DNase I (Thermo Scientific # EN0521). RNA was reverse transcribed into cDNA using iScript Reverse Transcription Supermix (Bio Rad #1708841). For qPCR, cDNA was amplified using gene specific primers (Supplementary Table 6) and PowerUp SYBR Green Master Mix (Applied Biosciences). RNA sequencing was performed at the London Regional Genomics Center. RNA-seq libraries were prepared and sequenced using NextSeq High 75 cycle kit. Sequencing reads were aligned to the human genome (Homo_sapiens.GRCh38.92) and read count files were generated using STAR aligner. We used DESeq2 to identify differentially expressed genes (53), and data displays were produced using BEAVR (54). RNA-seq read data is available through GEO226978.

### Gene set enrichment, network, clustering and over-representation analyses

For the CRISPR screen, GSEA of positively enriched (Log_2_ Fold Change > 1, p < 0.05) or negatively enriched (Log_2_ Fold Change < -1, p < 0.05) genes were identified using mageckGSEA (27). GO:BP pathways that were significantly enriched (p < 0.05, FDR < 0.05) were identified. For analysis of our EZ-resistance gene set, network analysis was performed on string-db.org (55), using an interaction score minimum of 0.400 and the following active interaction sources: Experiments, Databases, Co-expression, Neighbourhood, Gene Fusion and Co-occurrence. K-means clustering set to identify 3 clusters was used on the resulting network to identify clusters with significantly enriched gene ontology terms. For comparison of CRISPR screen and RNA-seq data, PANTHER overrepresentation test (FISHER test, FDR correction, Homo sapiens reference list) of significantly de-enriched genes (Log_2_ Fold Change < -1, p < 0.05) in T_f_ EZ vs. DMSO in the CRISPR screen analysis and significantly up-regulated genes (Log_2_ Fold Change > 1, p < 0.05) in the RNA-seq analysis was performed on geneontology.org (56, 57). GO:BP pathways that were significantly enriched (fold enrichment > 1.5, p value < 0.05, FDR < 0.05) were included.

### CBioPortal Prostate Tumour Data Analyses

For survival data, all prostate cancer studies within the CBioPortal database were selected and queried specifically for HOXA9 genomic amplification or gain using default settings (32). Analysis of indicated hormone sensitive prostate cancer studies (33, 58–60) including the TCGA firehose legacy (https://www.cancer.gov/tcga), and castration resistance (28, 61–63) prostate cancer datasets were also queried specifically for HOXA9 amplification or gain. For transcriptomic data from the SU2C/PCF Dream Team (28), FPKM PolyA mRNA data was used for analysis.

### Statistical Analyses

All statistical tests were performed as indicated in figure legends. For all analyses, results were considered statistically significant with *, ****p < 0.0001, ***p < 0.001, **p < 0.01, *p < 0.05, ND – Not Detected, ns – Not Significant

---

**Supplementary Figure 1.**
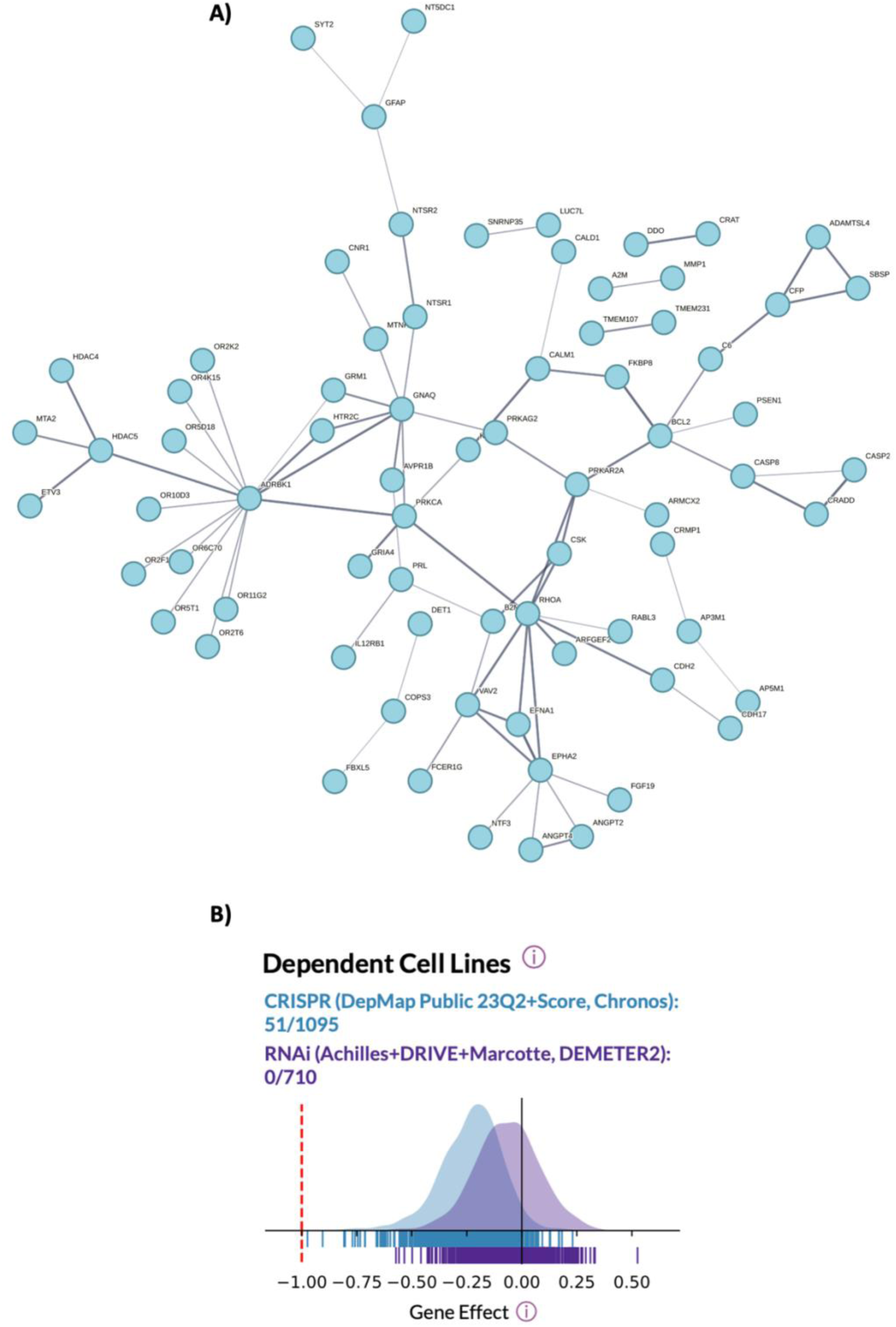
**A)** Cluster from network and clustering analysis of EZ resistance genes identified in CRISPR screen with central nodes including ADRBK1, RHOA and PRCKA **B)** Screenshot of distribution plot of gene effect scores for HOXA9 in all CRISPR (blue) and RNAi (purple) screens performed in cell lines reported in DepMap. The number of cell lines in which HOXA9 scored as a dependent gene in all CRISPR (blue) and RNAi (purple) screens is highlighted.

**Supplementary Figure 2.**
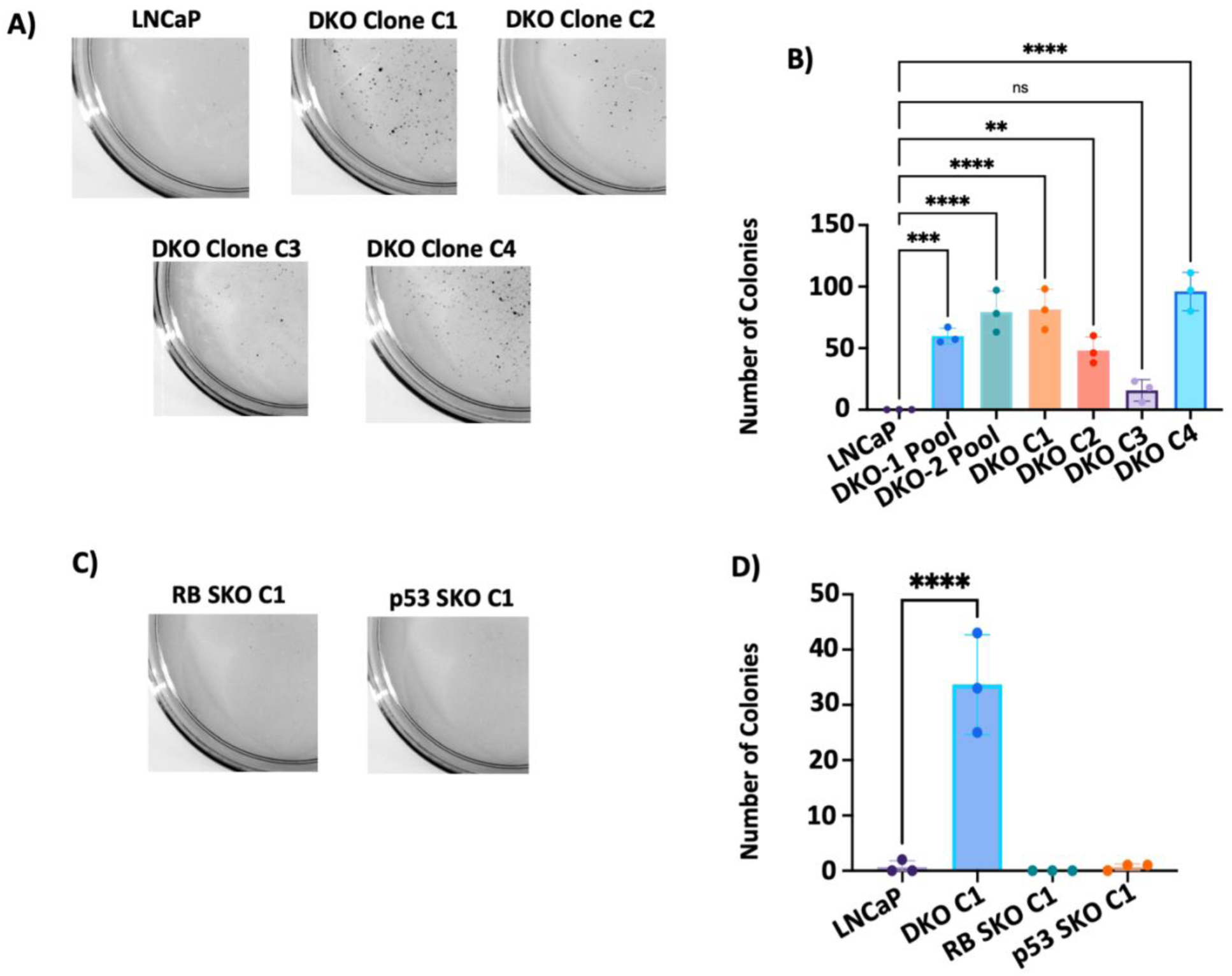
**A)** Representative images from colony forming assay of LNCaP WT and DKO clones. **B)** Analysis of colony forming assay from A. Number of colonies were counted using ImageJ software. **C)** Representative images from colony forming assay of RB SKO and p53 SKO clones. **D)** Analysis of colony forming assay from D including colony formation from LNCaP and DKO C1 cells.

**Supplementary Figure 3.**
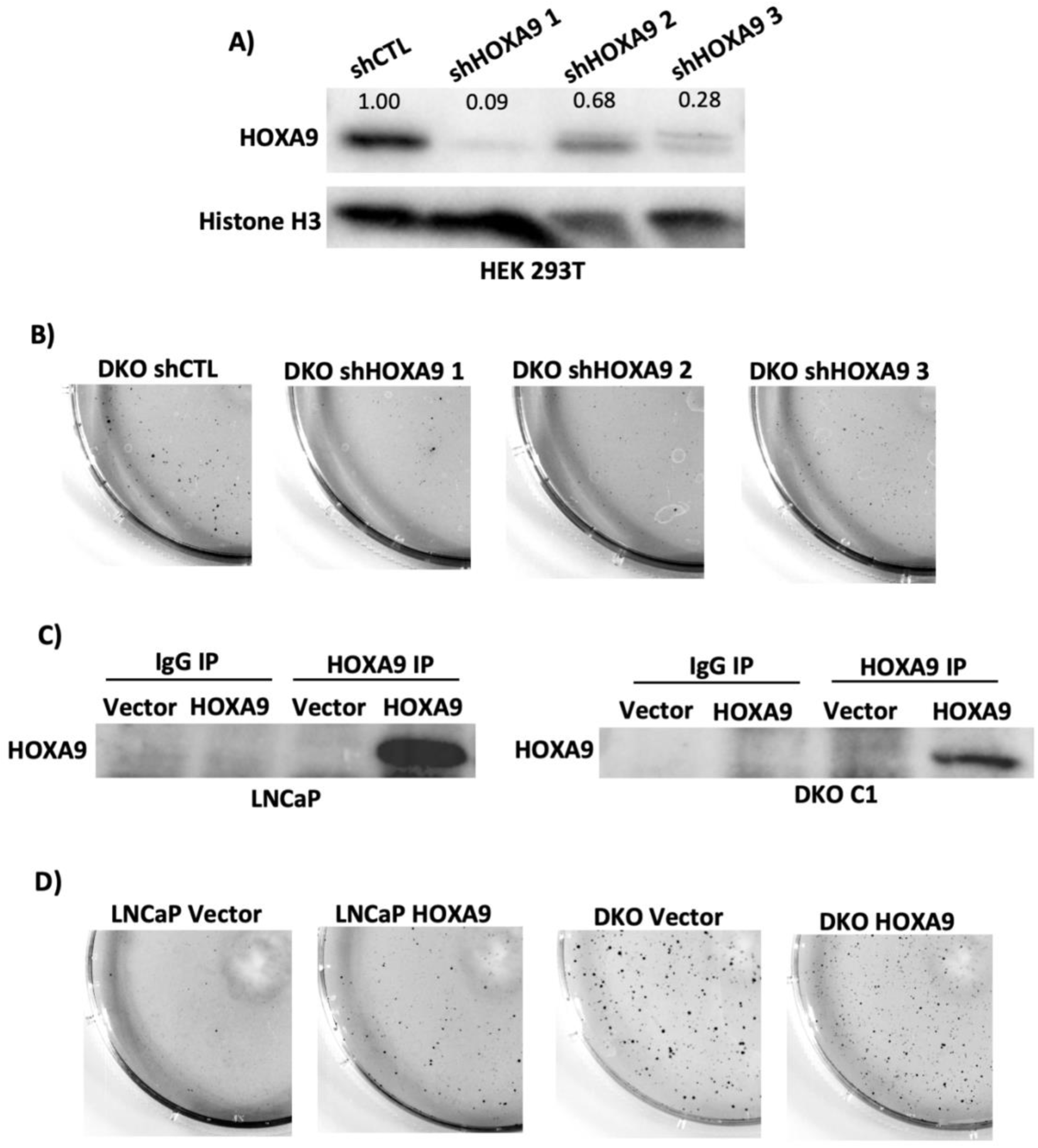
**A)** Representative western blot validating shRNA induced knockdown of HOXA9 in HEK-293T cells using 3 distinct shRNAs. Nuclear cell lysates from HEK 293T shCTL and shHOXA9 cells probed for HOXA9. Histone H3 is used as a loading control. Quantitation of band intensity for HOXA9 relative to Histone H3 loading control and normalized to shCTL cells is shown. **B)** Representative images from colony forming assay of RB-p53 DKO shCTL and DKO shHOXA9 clones. **C)** Western blot following immunoprecipitation of HOXA9 or IgG control from nuclear extracts of LNCaP and DKO empty vector (Vector)/HOXA9 overexpressing (HOXA9) cells and probed for HOXA9. **D)** Representative images from colony forming assay of LNCaP and DKO EV/HOXA9 OE cells.

